# SARS-CoV-2 Omicron-B.1.1.529 Variant leads to less severe disease than Pango B and Delta variants strains in a mouse model of severe COVID-19

**DOI:** 10.1101/2021.12.26.474085

**Authors:** Eleanor G. Bentley, Adam Kirby, Parul Sharma, Anja Kipar, Daniele F. Mega, Chloe Bramwell, Rebekah Penrice-Randal, Tessa Prince, Jonathan C. Brown, Jie Zhou, Gavin R. Screaton, Wendy S. Barclay, Andrew Owen, Julian A. Hiscox, James P. Stewart

**Author notes:** Corresponding author: E. mail.

## Abstract

COVID-19 is a spectrum of clinical symptoms in humans caused by infection with SARS-CoV-2. The B.1.1.529 Omicron variant is rapidly emerging and has been designated a Variant of Concern (VOC). The variant is highly transmissible and partially or fully evades a spectrum of neutralising antibodies due to a high number of substitutions in the spike glycoprotein. A major question is the relative severity of disease caused by the Omicron variant compared with previous and currently circulating variants of SARS-CoV-2. To address this, a mouse model of infection that recapitulates severe disease in humans, K18-hACE2 mice, were infected with either a Pango B, Delta or Omicron variant of SARS-CoV-2 and their relative pathogenesis compared. In contrast to mice infected with Pango B and Delta variant viruses, those infected with the Omicron variant had less severe clinical signs (weight loss), showed recovery and had a lower virus load in both the lower and upper respiratory tract. This is also reflected by less extensive inflammatory processes in the lungs. Although T cell epitopes may be conserved, the antigenic diversity of Omicron from previous variants would suggest that a change in vaccine may be required to mitigate against the higher transmissibility and global disease burden. However, the lead time to develop such a response may be too late to mitigate the spread and effects of Omicron. These animal model data suggest the clinical consequences of infection with the Omicron variant may be less severe but the higher transmissibility could still place huge burden upon healthcare systems even if a lower proportion of infected patients are hospitalised.

## Introduction

Since the emergence of SARS-CoV-2 in late 2019 several new variants of concern (VOC) have emerged and been associated with waves of infection across the globe. Most recently, the Delta VOC is estimated to have been responsible for over 99% of infections worldwide. The B.1.1.529 Omicron variant^1^ of SARS-CoV-2 that emerged recently in South Africa is causing considerable concern^2^. This variant appears to be highly transmissible and has many substitutions in the spike glycoprotein as well as elsewhere in the genome raising fears that the virus may escape from pre-existing immunity, whether acquired by vaccination or by prior infection^3^. Several mutations in the receptor binding domain and S2 region of the spike protein are predicted to impact transmissibility and affinity for the ACE-2 receptor^4^.

Infection of humans with SARS-CoV-2 results in a range of clinical courses, from asymptomatic to severe disease and subsequent death in at risk individuals but also a small proportion of otherwise healthy individuals across all age groups. Severe infection in humans is typified by cytokine storms ^5,6^, pneumonia, renlal failure and tissue specific immunopathological processes^7,8^. A small number of patients have no overt respiratory symptoms at all.

Animal models of COVID-19 present critical tools to fill knowledge gaps for the disease in humans. Compatibility with a more extensive longitudinal deep tissue sampling strategy and a controlled nature of infection are key advantages ^9^. Different animal species can be infected with wild-type SARS-CoV-2 to serve as models of COVID-19, these include mice, hamsters, ferrets^10^ and rhesus macaques and cynomolgus macaques^11^. The K18-hACE2 transgenic (K18-hACE2) mouse, where hACE2 expression is driven by the epithelial cell cytokeratin-18 (K18) promoter, was developed to study SARS-CoV pathogenesis^12^. This mouse is now widely used as a model that mirrors many features of severe COVID-19 infection in humans to develop understanding of the mechanistic basis of lung disease and to test pharmacological interventions^13,14^.

With the apparent high transmissibility of the Omicron variant as well as the ability to evade pre-existing immunity, we sought to rapidly assess the relative pathogenicity against previous isolates of SARS-CoV-2. To do this, K18-hACE2 mice were infected with a Pango B lineage variant, Delta and Omicron variants of SARS-CoV-2 and their relative pathogenesis and viral loads compared.

## Results

### Infection of hACE2 mice with Omicron variant leads to less severe clinical signs and recovery

To assess the relative pathogenicity of the Omicron variant, the established K18-hACE2 mouse model of SARS-CoV-2 was utilised^12^. A near clinical (B.1.1.529) Omicron variant isolate from the UK^3^ was used along with two variants of known provenance of SARS-CoV-2 as comparators. These were representative of a variant from the initial outbreak in the UK (strain hCoV-19/England/Liverpool_REMRQ0001/2020; Pango B)^15^ and a Delta variant (B1.617.2). Sequencing of the virus stocks in house demonstrated that the stock viruses did not contain unexpected deletions/insertions or substitutions and matched non-synonymous and synonymous signatures associated with each variant. A schematic of the experimental design is shown in Fig. 1.

**Figure 1:**
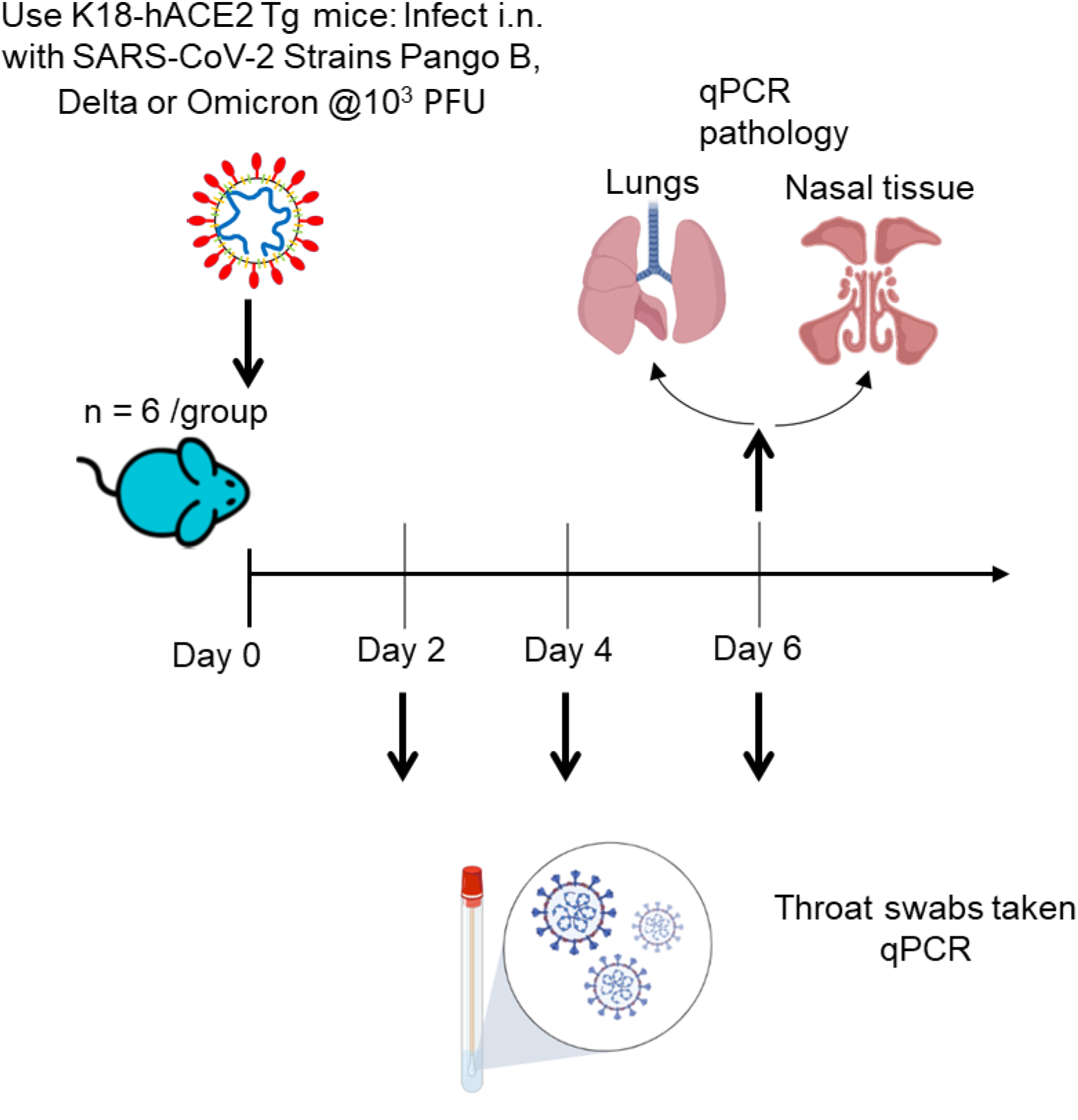
Schematic diagram of the experimental design for infection of K18-hACE2 mice with Pango B, Delta and Omicron variants of SARS-CoV-2. Infection of these mice with SARS-CoV-2 provides a model that resembles severe infection in humans that had not been vaccinated or treated with currently available licenced therapeutics. The major measure of disease was weight loss followed by end of experiment pathology. Throat swabs were taken daily for RT-qPCR to determine and compare viral loads.

Three groups of mice (n = 6 per group) were used and inoculated intranasally with 10^3^ PFU of each SARS-CoV-2 variant. The results (Fig. 2) indicated that mice infected with the Pango B and Delta variants displayed an expected pattern of weight loss^16^. This included a substantial drop in weight at day 3 which progressed in Pango B and Delta variant infected mice until euthanasia at day 6 post-infection (mean 20% loss). Mice infected with the Omicron variant had a similar degree and pattern of weight loss up to day 5 post-infection (mean 16% loss). However, at day 6 post-infection, they displayed a statistically significant recovery in weight loss as compared to the other groups (mean 10% loss; *p* < 0.05 2-way ANOVA with Bonferroni post-test).

**Figure 2:**
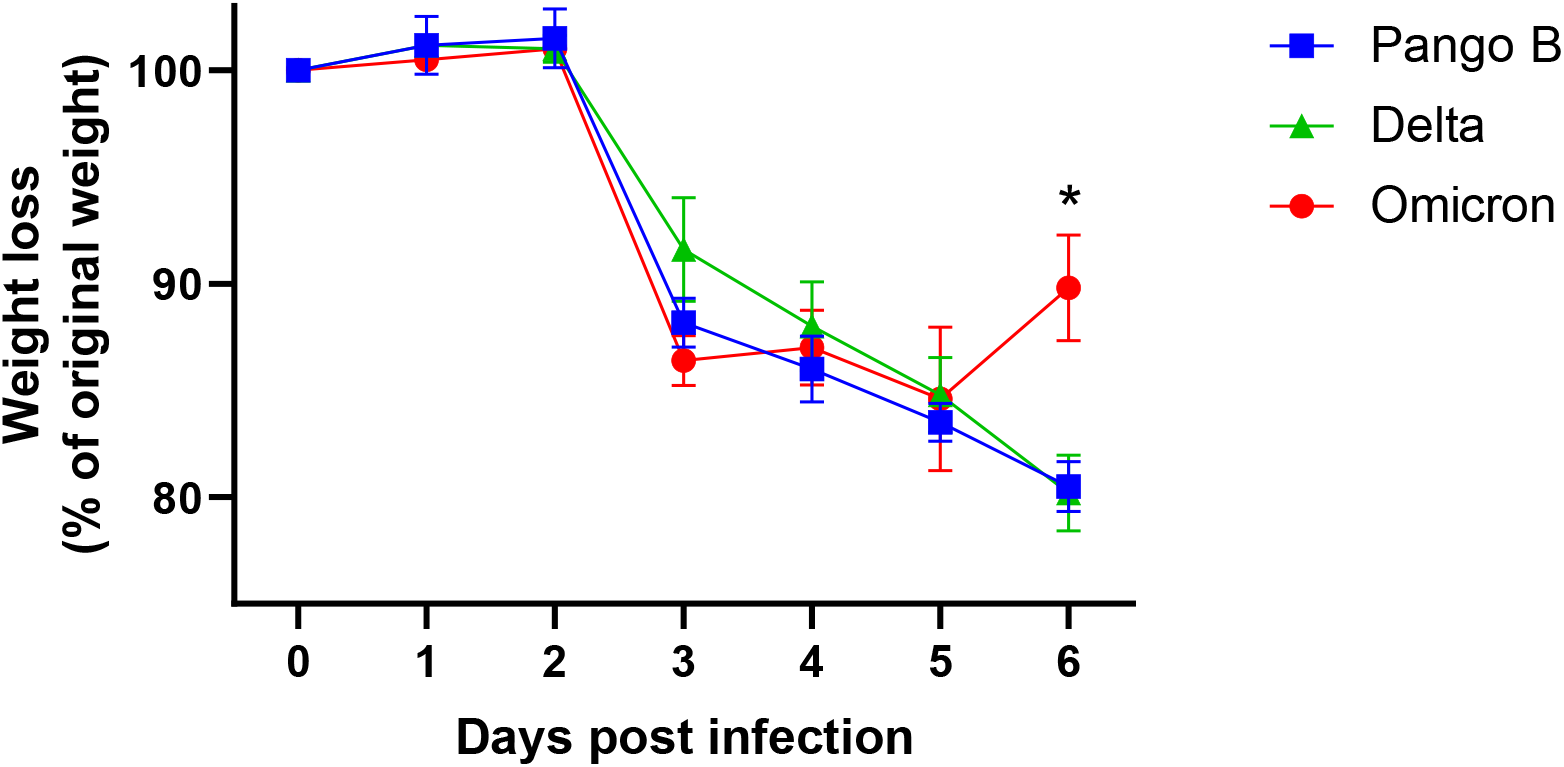
Weight loss after infection with SARS-CoV-2 variants. K18-hACE2 mice were challenged intranasally with 10^3^ PFU SARS-CoV-2 variants as indicated. Mice were monitored for weight loss at indicated time-points. (n = 6). Data represent the mean value ± SEM. Comparisons were made using a repeated-measures two-way ANOVA (Bonferroni post-test). *Represents *p* < 0.05

### Mice infected with Omicron variant have lower viral load

To determine the viral load in animals infected with each variant, total RNA was extracted from oral swabs taken at days 2, 4, and 6 post-infection. At the experimental end point lungs and nasal tissue were also taken. Viral loads were quantified using qRT-PCR to measure viral RNA as a proxy. At day 2 post infection, mice infected with Omicron variant exhibited 100-fold lower levels of viral RNA than Pango-B and Delta variant-infected animals (mean 2.5 × 10^3^ vs 1 × 10^5^ and 6 × 10^4^ copies of N/μg of RNA respectively). The difference between Omicron and Pango B infected mice was significant (*p* < 0.05) (Fig, 3A). At day 4 post-infection, the viral loads were similar between all groups of mice. At day 6 post-infection, the viral load was approximately 10-fold higher in the Pango B and Delta variant infected mice than the Omicron-infected mice and this difference was significant between Delta and Omicron infected animals (Fig. 3A). At this stage, viral loads in nasal tissue were significantly lower in Omicron-infected mice than in the other two groups (approx. 100 fold) (Fig. 3B). Similary, viral loads in the lungs were significantly lower (approx. 100 fold) in Omicron versus Pango B and Delta variant-infected mice (Fig. 3C).

**Figure 3:**
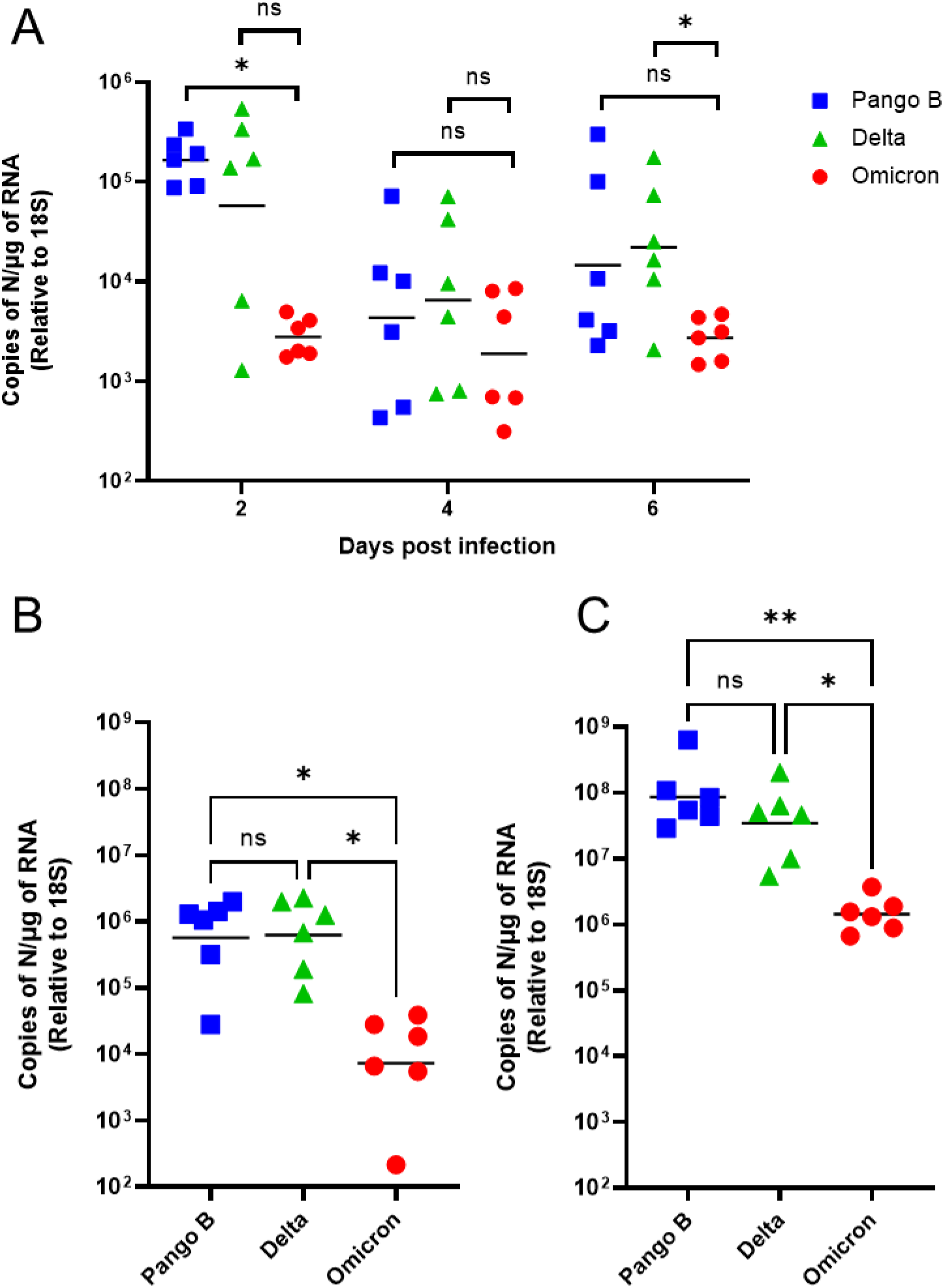
SARS-CoV-2 loads in infected mice. K18-hACE2 mice were challenged intranasally with 10^3^ PFU SARS-CoV-2 Pango B, Delta or Omicron variants (n = 6 per group). RNA extracted from oral swabs **(A)** nasal tissue **(B)** and lungs **(C)** was analysed for virus levels by qRT-PCR for the N gene. Assays were normalised relative to levels of 18S RNA. Data for individual animals are shown with the geometic mean value represented by a black line Comparisons were made using the Mann-Whitney U test (A) and the Kruskal-Wallis test with Dunn’s multiple comparisons (B and C). * represents *p* < 0.05; ** represents *p* < 0.01

### Mice infected with Omicron variant exhibit less severe pneumonia

To determine the extent of inflammatory changes in the lungs, the left lobes were examined grossly, by histology and by immunohistology for the detection of SARS-CoV-2 nucleoprotein expression. In animals infected with the Pango B or Delta variants, the lung showed large patchy areas of dark red discoloration (Fig. 4). Histologically, a mild multifocal to diffuse increase in interstitial cellularity was seen. This was accompanied by partly extensive areas where alveoli contained a few desquamed alveolar macrophages/type II pneumocytes, occasional degenerate cells as well as neutrophils, lymphocytes and macrophages. There were also areas where alveoli exhibited activated type II pneumocytes and occasional syncytial cells. There was extensive viral antigen expression in type I and II pneumocytes in both unaltered alveoli and those involved in the inflammatory processes. In the latter, macrophages were also found to be positive (Fig. 4). In addition, mild to moderate patchy to circular periarterial lymphocyte and macrophage dominated mononuclear infiltration, mild arteritis and vascular endothelial cell activation was also observed (Fig. 4), as previously described^16–18^. The lungs of Omicron variant infected mice appeared grossly widely unaltered (Fig. 4). Similarly, the histological examination revealed widely unaltered parenchyma, with focal mild increase in interstitial cellularity, small, more loosely consolidated areas (Fig. 4) with activated type II pneumocytes, occasional syncytial cells and degenerate cells, some desquamed macrophages/type II pneumocytes in alveolar lumina, and some infiltrating macrophages and neutrophils. A few small patchy leukocyte aggregates (macrophages, neutrophils, lymphocytes) were occasionally observed adjacent to small arteries. Viral antigen expression was detected in 5 of the 6 animals, in type I and II pneumocytes and in macrophages in focal inflammatory processes (Fig. 4). A few arteries exhibited a mild vasculitis and/or mild patchy to circular mononuclear perivascular infiltrates.

**Figure 4:**
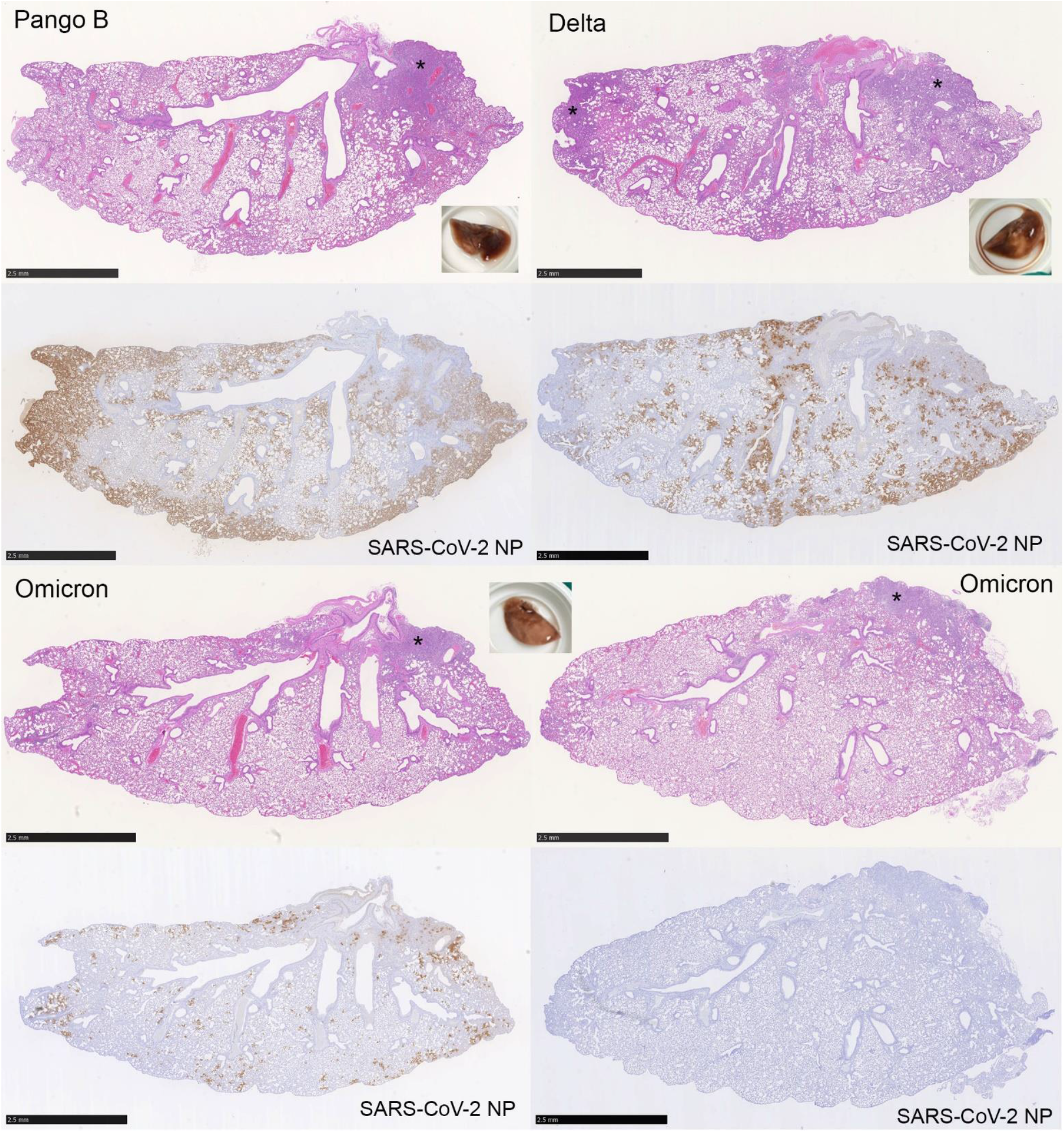
Gross and histological features and viral antigen expression in the lungs of SARS-CoV-2 infected mice. K18-hACE2 mice were challenged intranasally with 10^3^ PFU SARS-CoV-2 Pango B, Delta or Omicron variants (n = 6 per group). Gross images and longitudinal sections of the left lung, stained with HE (top) and for SARS-CoV-2 nucleoprotein (NP) expression (bottom). **Pango B and Delta infected animals.** Gross examples of the lungs, with large patches of dark red discolouration. Histologically, there are large areas of consolidation (*). Overall, the lung parenchyma exhibits an increased interstitial cellularity. HE stain. Immunohistology shows extensive viral antigen expression throughout the lung parenchyma. **Omicron infected animals**. Gross example of the lungs which appeared unaltered. Histologically, the lung parenchyma appears widely unaltered. There are only small patches of consolidation (*). Immunohistology shows multiple small patches of alveoli with viral antigen expression in one animal (left), whereas viral antigen cannot be detected in another mouse in the same group (right).

## Discussion

In this study, infection with the SARS-CoV-2 Omicron variant led to less severe weight loss, lower viral loads in oral swabs, lungs and nasal tissue and less severe pathological changes in lung tissue than animals infected with either Pango B or Delta variant viruses. Weight loss is the best objective measure of clinical severity in mouse models of COVID-19^16^. Our results show that while Omicron variant-infected mice lose weight initially as fast as when infected with Pango B and Delta variant-infected animals, they recover significantly between day 5 and 6 post-infection. This is also reflected by less extensive inflammatory processes and less virus antigen in the lungs. These results indicate that the Omicron variant causes a disease which is less clinically severe than that caused by Pango B or Delta variant viruses.

Mice infected with the Omicron variant had significantly lower viral loads as measured by RT-qPCR than those in Pango B or Delta variant-infected animals. This strongly suggests lower levels of virus replication in Omicron-infected mice in both the upper and lower respiratory tract. A relative decrease in viral replication and hence load fits with the observed decrease in clinical and pathological severity and recovery seen in Omicron-infected mice. Previous data from animal models suggested a relationship between viral load and disease severity^10^.

No animal model can predict with absolute certainty the consequences of infection in humans. However, the data presented here strongly suggest that the clinical consequences of infection in humans with the Omicron variant may be less profound than infection with either the Delta variant or the original Pango B viruses. While it is still very early in the spread of the Omicron variant, a decrease in clinical severity for Omicron-infected patients is starting to be reported, which is supported by the observations in this study.

The Omicron variant would appear to be easily transmitted, and indeed is spreading at a faster rate than Delta which has been the dominant variant worldwide for much of 2021. One factor in efficiency of spread and transmission, apart from immune status, is the level of replication and viral loads, especially in the upper respiratory tract. The data presented here indicate that, at least in this mouse model, there are lower viral loads of the Omicron variant in both the lower and upper respiratory tract than the other variants. If this is the case in humans, the reasons for the high efficiency of transmission of the Omicron variant must lie in other factors such as efficiency of binding to host cells, or in evading the initial host defences in contacts, and/or behavioural patterns.

Our results, and emerging from human observational studies, suggest that the Omicron variant may lead to less severe and/or more rapid recovery from clinical disease reflected in reduced hospitalisation. However, the reported high transmissibility and the fact that the Omicron variant will evade much of the pre-existing immunity in populations as well as existing antibody-based therapies argues for the maintenance of other measures such as social distancing, mask-wearing and restriction of indoor contacts to obviate a potentially catastrophic impact on healthcare saturation.

## Acknowledgements

This work was funded by MRC grant MR/W005611/1, G2P-UK; A National Virology Consortium to address phenotypic consequences of SARS-CoV-2 genomic variation (JPS, JAH, WSB) and by the U.S. Food and Drug Administration Medical Countermeasures Initiative contract (75F40120C00085) ‘Characterization of severe coronavirus infection in humans and model systems for medical countermeasure development and evaluation’ (JAH). The article reflects the views of the authors and does not represent the views or policies of the FDA. The authors are grateful to Sabina Wunderlin, Histology Laboratory, Institute of Veterinary Pathology, Vetsuisse Faculty, University of Zurich, for excellent technical support.

## Methods

### Viruses

A UK strain of SARS-CoV-2 (hCoV-2/human/Liverpool/REMRQ0001/2020), Pango lineage B, was cultured from a nasopharyngeal swab from a patient^15^. The sequence was submitted to Genbank, accession number MW041156.

The B.1.617.2 (delta variant) hCoV-19/England/SHEF-10E8F3B/2021 (GISAID accession number EPI_ISL_1731019), was kindly provided by Prof. Wendy Barclay, Imperial College London, London, UK through the Genotype-to-Phenotype National Virology Consortium (G2P-UK). Sequencing confirmed it contained the spike protein mutations T19R, K77R, G142D, Δ156-157/R158G, A222V, L452R, T478K, D614G, P681R, D950N.

The B.1.1.529/BA.1 (Omicron variant) isolate M21021166 was originally isolated by Prof Gavin Screaton, University of Oxford^3^, UK and then obtained from Prof. Wendy Barclay, Imperial College London, London, UK through the Genotype-to-Phenotype National Virology Consortium (G2P-UK). Sequencing confirmed it contained the spike protein mutations A67V, Δ69-70, T95I, G142D/Δ143-145, Δ211/L212I, ins214EPE, G339D, S371L, S373P, S375F, K417N, N440K, G446S, S477N, T478K, E484A, Q493R, G496S, Q498R, N501Y, Y505H, T547K, D614G, H655Y, N679K, P681H, N764K, A701V, D796Y, N856K, Q954H, N969K, L981F.

The titres of all isolates were confirmed on Vero E6 cells and the sequences of all stocks confirmed.

### Biosafety

All work was performed in accordance with risk assessments and standard operating procedures approved by the University of Liverpool Biohazards Sub-Committee and by the UK Health and Safety Executive. Work with SARS-CoV-2 was performed at containment level 3 by personnel equipped with respirator airstream units with filtered air supply.

### Mice

Animal work was approved by the local University of Liverpool Animal Welfare and Ethical Review Body and performed under UK Home Office Project Licence PP4715265. Mice carrying the human ACE2 gene under the control of the keratin 18 promoter (K18-hACE2; formally B6.Cg-Tg(K18-ACE2)2Prlmn/J) were purchased from

Charles River. Mice were maintained under SPF barrier conditions in individually ventilated cages.

### Virus infection

Animals were randomly assigned into three cohorts. For SARS-CoV-2 infection, mice were anaesthetized lightly with isoflurane and inoculated intra-nasally with 50 μl containing 10^3^ PFU SARS-CoV-2 in PBS. They were sacrificed at variable time-points after infection by an overdose of pentabarbitone. Animals were dissected and tissues collected immediately for downstream processing.

### RNA extraction and DNase treatment

The upper lobes of the right lung were homogenised in 1ml of TRIzol reagent (Thermofisher) using a Bead Ruptor 24 (Omni International) at 2 m/s for 30 s. The homogenates were clarified by centrifugation at 12,000xg for 5 min before full RNA extraction was carried out according to manufacturer’s instructions. RNA was quantified and quality assessed using a Nanodrop (Thermofisher) before a total of 1 μg was DNase treated using the TURBO DNA-free™ Kit (Thermofisher) as per manufacturer’s instructions.

### qRT-PCR for viral load

Viral loads were quantified using the GoTaq® Probe 1-Step RT-qPCR System (Promega). For quantification of SARS-COV-2 the nCOV_N1 primer/probe mix from the SARS-CoV-2 (2019-nCoV) CDC qPCR Probe Assay (IDT) were utilised while the standard curve was generated via 10-fold serial dilution of the 2019-nCoV_N_Positive Control (IDT) from 10^6^ to 0.1 copies/reaction. Murine 18S primers and probe sequences were utilised at 400nM and 200nM respectively. The 18s standard was generated by the amplification of a fragment of the murine 18S cDNA using the primers F: ACCTGGTTGATCCTGCCAGGTAGC and R: GCATGCCAGAGTCTCGTTCG. cDNA was generated using the SuperScript IV reverse transcriptase kit (Thermofisher) and PCR carried out using Q5® High-Fidelity 2X Master Mix (New England Biolabs) as per manufacturer’s instructions. Both PCR products were purified using the QIAquick PCR Purification Kit (Qiagen) and serially diluted 10-fold from 10^10^ to 10^4^ copies/reaction to form the standard curve.

### Histology and immunohistology

The left lungs were fixed in 10% buffered formalin for 48 h and stored in 70% ethanol until trimming for histological examination and routinely paraffin wax embedding. Consecutive sections (3–5 μm) were prepared and routinely stained with hematoxylin-eosin (HE) or subjected to immunohistochemistry (IHC) for the detection of SARS-CoV-2 antigen as previously described^16^. IHC was performed in an autostainer (Agilent) using a rabbit polyclonal anti-SARS-CoV nucleoprotein antibody (Rockland, 200-402-A50) and the horseradish peroxidase (HRP) method. Briefly, sections were deparaffinized and rehydrated through graded alcohol. Antigen retrieval was achieved by 20 min incubation in citrate buffer (pH 6.0) at 98 °C in a pressure cooker. This was followed by incubation with the primary antibody (diluted 1:6,000 in dilution buffer; Dako) overnight at 4 °C, a 10 min incubation at RT with peroxidase blocking buffer (Agilent) and a 30 min incubation at RT with Envision+System HRP Rabbit (Agilent). The reaction was visualized with diaminobenzidin (DAB; Dako) for 10 min at RT. After counterstaining with hematoxylin for 2 s, sections were dehydrated and coverslipped.

